# HIF2α negatively regulates MYCN protein levels and promotes a low-risk noradrenergic phenotype in neuroblastoma

**DOI:** 10.1101/2024.11.20.624001

**Authors:** Juan Yuan, Subhamita Maitra, Eirini Antoniou, Jiacheng Zhu, Wenyu Li, Ilknur Safak Demirel, Kostantinos Toskas, Iria Laura Martinez, Lacin Ozcimen, Henrik Lindehell, Per Kogner, Oscar C. Bedoya-Reina, Susanne Schlisio, Johan Holmberg

## Abstract

The role of HIF2α, encoded by *EPAS1*, in neuroblastoma remains controversial. Here we demonstrate that induction of high levels of HIF2α in MYCN-amplified neuroblastoma cells results in a rapid and profound reduction of the oncoprotein MYCN. This is followed by an upregulation of genes characteristic of noradrenergic cells in the adrenal medulla. Additionally, upon induction of HIF2α, the proliferation rate drops substantially, and cells develop elongated neurite-like protrusions, indicative of differentiation. *In vivo* HIF2α induction in established xenografts significantly attenuates tumour growth. Notably, analysis of sequenced neuroblastoma patient samples, revealed a negative correlation between *EPAS1* and *MYCN* expression and a strong positive correlation between *EPAS1* expression, high expression levels of noradrenergic markers, and improved patient outcome. This was paralleled by analysis of human adrenal medulla datasets wherein *EPAS1* expression was prominent in populations with high expression levels of genes characteristic of noradrenergic chromaffin cells. Our findings show that high levels of HIF2α in neuroblastoma, leads to drastically reduced MYCN protein levels, cell cycle exit, and noradrenergic cell differentiation. Taken together, our results challenge the dogma that HIF2α acts as an oncogene in neuroblastoma and rather suggest that HIF2α has potential tumour suppressor capacity in this particular disease.

**Significance statement:** HIF2α has been proposed as a neuroblastoma oncogene and a tractable target for clinical intervention, this has been questioned by several studies. Thus, it is necessary to move beyond correlative studies and better determine the function of HIF2α in neuroblastoma. Our study shows that induced expression of HIF2α in *MYCN-*amplified neuroblastoma substantially reduces MYCN protein levels and attenuates proliferation while it induces several features of noradrenergic differentiation and impedes xenograft tumour growth. Together with bioinformatic analysis of sequenced neuroblastoma patient samples and the developing human adrenal medulla, this couples HIF2α to low-risk neuroblastoma with a substantially better patient outcome. Thus, in neuroblastoma HIF2α exhibit tumour suppressor capacity rather than oncogenic capacity.

## Introduction

Neuroblastoma arises in the sympathoadrenal lineage and is the most frequent extracranial solid childhood cancer, with a high degree of clinical heterogeneity ranging from spontaneous regression to fatal progression (1). Like in many paediatric malignancies, the frequency of recurring somatic mutations is low. Instead, the disease is characterized by loss and gain of distinct chromosomal regions, for example loss of 1p36, loss of 11q and gain of 17q. The best characterized neuroblastoma oncogene is *MYCN* which is amplified in roughly 20% of high-risk cases, of which less than 50% survive (2). Even though MYCN has a short protein half-life, the *MYCN* amplification generates extremely high levels of MYCN protein (3). Targeting of MYCN with specific inhibitors has proven challenging, largely due to the intrinsically disorganised structure and lack of a suitable binding pocket. Given the central oncogenic role of MYCN, any process that can significantly reduce MYCN protein levels in an *MYCN*-amplified neuroblastoma would most probably also reduce tumour growth and aggressiveness.

In certain other types of tumours, e.g., subtypes of renal clear cell carcinoma and cases of sporadic paraganglioma, HIF2α is a well-studied oncogenic driver of the disease (4) and a similar role for HIF2α has been suggested in neuroblastoma (5). However, such an oncogenic role for HIF2α has been challenged in several studies (6–10). Despite several hundred bulk RNA-sequenced/arrayed neuroblastoma tumours exhibiting a highly significant correlation between high levels of *EPAS1* expression and low-risk tumours with favourable outcome (7), some studies have continued to argue that HIF2α acts to fuel tumour growth, partly by imposing a an immature state with stem cell-like features that blocks differentiation (5, 11–15). However, experimental evidence supporting an oncogenic role for HIF2α in neuroblastoma remains ambiguous.

The generation of neuroblastoma has been tightly linked to the developmental context of the sympathoadrenal system and advances in single-cell sequencing have provided a much more detailed understanding of this lineage. A series of studies has mapped the developmental trajectory of sympathoadrenal cells in both mice and humans (16–23). Importantly, these studies include efforts to align expression profiles of developmental stages with neuroblastoma, which have revealed that neuroblastoma substantially resembles the sympathoblast/neuroblast lineage (16, 20, 21, 23, 24). In contrast, cells within the chromaffin lineage exhibit significantly less features of high-risk neuroblastoma (20, 21).

Here we show that *EPAS1* expression in the developing sympathoadrenal system is correlated with key genes required for noradrenergic cell differentiation and function. This reflects previous studies showing that basal levels of HIF2α are required for the normal development of the catecholaminergic phenotype of sympathoadrenal cells, including the expression of enzymes necessary to produce noradrenaline (25). Overexpression of HIF2α in *MYCN*-amplified neuroblastoma cells rapidly induces expression of a cohort of genes also enriched in the chromaffin lineage during normal adrenal gland development, including key enzymes such as *TH, DDC* and *DBH,* but not *PNMT,* indicating a bias towards noradrenergic chromaffin cells (N-cells) rather than adrenergic E-type cells. This is accompanied by a significant reduction in cellular proliferation. Importantly, levels of the MYCN oncoprotein are rapidly diminished, resulting in decreased expression of MYCN target genes. In addition, induction of HIF2α in already established neuroblastoma xenografts significantly impedes tumour growth. In sum, our data challenge the concept of HIF2α as a neuroblastoma oncogene and show that HIF2α is rather associated with decreased MYCN levels and low-risk tumours as well as noradrenergic differentiation.

## Results

### *EPAS1* expression is positively correlated with genes highly expressed in noradrenergic chromaffin cells and in low-risk neuroblastoma

Through analysis of data from Furlan et al. (18) we previously mapped *EPAS1* expression to the chromaffin lineage during mouse adrenal medulla development (7). To understand whether *EPAS1* expression is associated with the chromaffin lineage also during human development, we utilized previously published data from Jansky et al. (20) (Fig. 1D). We plotted the expression of *EPAS1* on to the single cell RNAseq data set derived from human embryonic adrenal medullas, ranging from 7 to 17 weeks post conception (PCW). High levels of *EPAS1* were evident in two populations designated as connecting progenitor cells and as chromaffin cells (Fig. 1 A). Expression of dopa decarboxylase (*DDC*), an enzyme converting L-DOPA to dopamine, which in turn is a necessary step to produce noradrenaline, is also enriched in these two populations (Fig. 1B), as are other key enzymes in this pathway, *TH* and *DBH* (S1A-B). In addition, monoamine transporters *SLC18A1* and *SLC18A2*, necessary for adrenal monoamine homeostasis as well as the cell cycle inhibitor *CDKN1C* and the tumour suppressor *NDRG1*, are also enriched for in the same populations (Fig. 1C and Fig. S1C-D). There is however no correlation between *EPAS1* and *PNMT*, the enzyme necessary to produce adrenaline from noradrenaline (Fig. S1E). This implies that *EPAS1* in human adrenal medulla development is associated with the N-type of chromaffin cells but to a lesser degree with the adrenaline producing E-type chromaffin cells.

**Figure 1.**
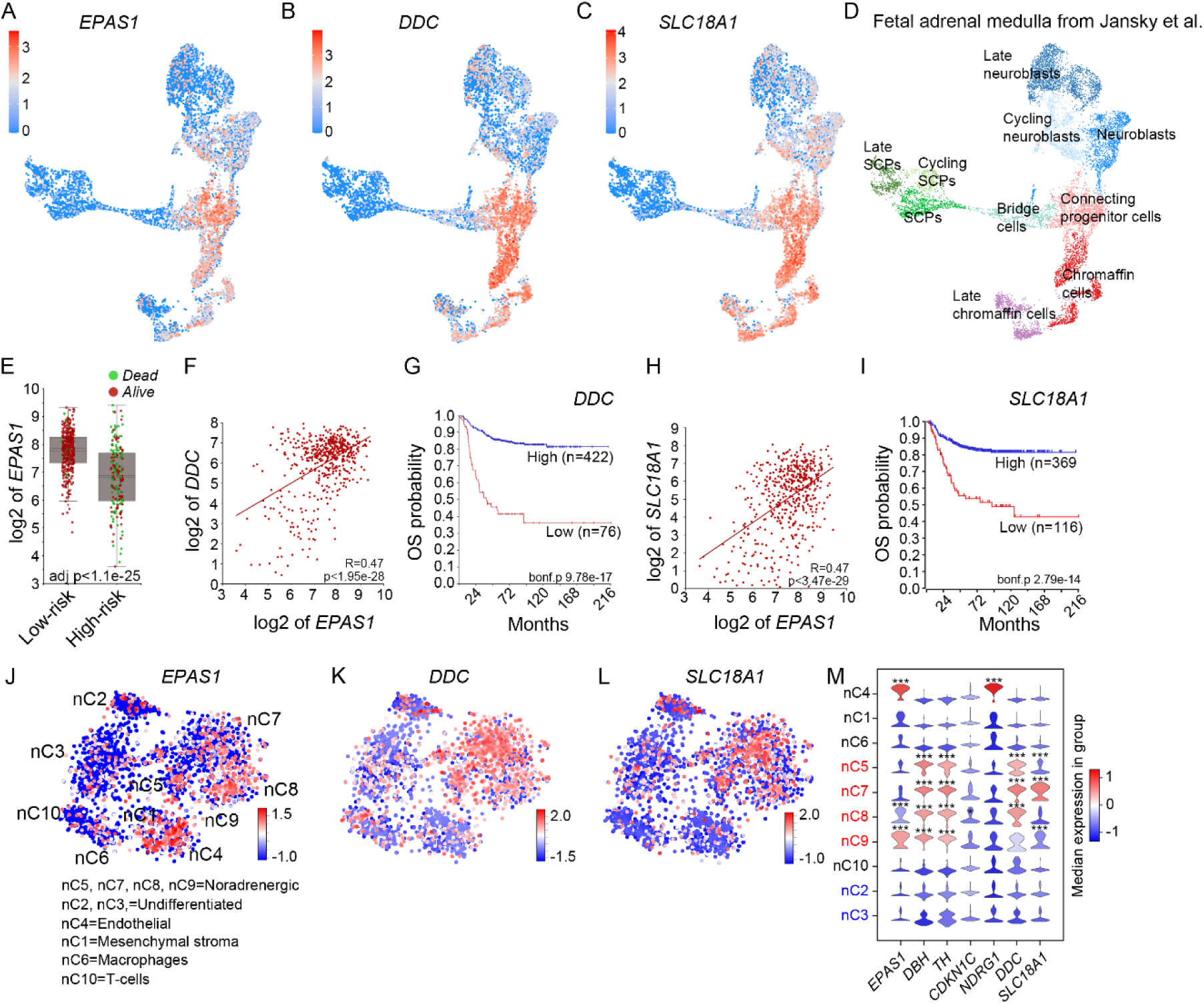
*EPAS1* expression is associated with the chromaffin cell lineage and low-risk neuroblastoma. (A-D) Visualization of *EPAS1* (A), *DDC* (B) and *SLC18A1* (C) expression in UMAP plot of the developing human adrenal medulla 7-17 PCW (D), dataset from Jansky et al. (20). (E) In neuroblastoma tumors *EPAS1* expression is positively correlated with low-risk tumors lacking *MYCN* amplification. (F) Expression of *DDC* and *EPAS1* are positively correlated in neuroblastoma tumors. (G) *DDC* expression is correlated to increased overall survival. (H) Expression of *SLC18A1* and *EPAS1* are positively correlated in neuroblastoma tumors. (I) *SLC18A1* expression is correlated to increased overall survival. (J) Mapping of *EPAS1* expression in tSNE of neuroblastoma single cell nuclei show enrichment in endothelial cells, and in noradrenergic low-risk tumor cells, dataset from Bedoya-Reina et al. (16). (K) *DDC* expression is enriched for in noradrenergic low-risk tumor cells. (L) *SLC18A1* expression is enriched for in noradrenergic low-risk tumor cells. (M) Violin plots depicting median expression of the indicated genes in each group. Groups in red font represent noradrenergic neuroblastoma and groups in blue font represent undifferentiated high-risk neuroblastoma. *** Indicates FDR<0.01.

Recent studies show that, in neuroblastoma, the majority of cells predominantly resemble sympathoblasts/neuroblasts, rather than chromaffin cells. This is most clearly evident in high-risk tumours (20, 21, 26). To understand whether *EPAS1* expression also is correlated with chromaffin markers in neuroblastoma tumours we analyzed the R2 database, utilizing the 498SEQC dataset (27, 28). Expectedly, *EPAS1* expression was significantly higher in the low-risk tumours (Fig. 1E). Genes defining the noradrenergic phenotype, for example, the key rate limiting enzymes for the production of monoamines *TH*, *DDC* and *DBH* as well as the monoamine transporters *SLC18A1* and *SLC18A2* (29) correlate strongly with *EPAS1* expression. In addition, just as for *EPAS1*, high levels of expression of all these genes are strong predictors of improved patient outcome (Fig. 1 F-I and Fig. S1 F-I, K-N). Thus, in combination with previous studies showing that HIF2α is necessary for the catecholaminergic phenotype in sympathoadrenal cells (25) and in the embryonic organ of Zuckerkandl (30), this suggests that high levels of *EPAS1* expression is associated with noradrenergic chromaffin differentiation and that in other cell types of the adrenal medulla the expression levels are generally low. There is a similar lack of overlap between *PNMT* and *EPAS1* expression in neuroblastoma as in the developing adrenal medulla (Fig. S1J), potentially reflecting a N-type like, rather than E-type like, chromaffin identity in cells with elevated *EPAS1* levels.

Through analysis of several bulk sequenced data sets of neuroblastoma patient samples we have previously shown that high expression levels of *EPAS1* is strongly correlated with increased patient survival and features typical of low-risk tumours (6, 7, 31). To understand whether this also could be reflected in single cell sequenced neuroblastoma, we inspected 11 previously single cell sequenced tumours (16). This revealed that *EPAS1*, besides from a strong enrichment in endothelial cells, is significantly enriched in clusters nC8 and nC9 from noradrenergic low-risk tumours (NOR-clusters) characterized by high expression of noradrenergic markers such as *DDC* and *SLC18A1* and significantly negatively correlated with undifferentiated high-risk neuroblastoma with high *MYCN* expression (Fig. 1 J-M and Fig. S1 O-R) (16). Thus, *EPAS1* is associated with a more differentiated noradrenergic cellular state. In contrast, neuroblastoma cells belonging to more undifferentiated high-risk tumours exhibit lower levels of *EPAS1* (Fig. 1J and M). To further validate this pattern, we utilized a recently published study wherein single cell/nuclei sequencing data from 6 different studies comprising of 68 samples from 61 different neuroblastoma patients had been combined (24). In the “NBAtlas” tool provided in this publication we mapped the expression of *EPAS1*, *DDC* and *MYCN* in high-and low-risk neuroblastoma. Supporting our previous analysis, there is an overlap of *EPAS1* and *DDC* expression, predominantly in low-risk tumour cells whereas the *MYCN* transcript is strongly enriched for in the high-risk tumour cells (Fig. S2A-D).

To map the expression of *EPAS1* in neuroblastoma *in situ,* we performed RNA-scope with probes for *EPAS1*, *TH* and the endothelial marker *ENG* (*Endoglin*) in tumours of different stages. In low-risk stage 4S and 2B tumours *EPAS1* co-localized with *ENG* but there were also several areas with strong co-localization between *EPAS1* and *TH* where *ENG* was not present (Fig. S3 A-B). In a high-risk stage 4 tumour, *EPAS1* was typically only expressed in blood vessels expressing high levels of *ENG.* Interestingly, in a restricted area of the tumour containing strong focal expression of *TH* there was also *EPAS1* expression without *ENG*, indicating that also in high-risk tumours there are restricted regions with co-expression of genes usually associated with low-risk tumours (Fig. S3C).

### High levels of HIF2α promote reduced proliferation and features of noradrenergic chromaffin differentiation in *MYCN-*amplified neuroblastoma cells

The clear correlation of *EPAS1* with increased patient survival, low-risk tumours and noradrenergic differentiation prompted us to test if increased levels of HIF2α in neuroblastoma cells would affect key cellular properties such as proliferation and differentiation. To address this, we utilized the *piggyBac* transposon system (32) to overexpress a stabilized version of HIF2α (33) under the control of doxycycline (*iHIF2α*) (Fig. 2A). In *MYCN*-amplified LAN-1 neuroblastoma cells induction of *iHIF2α* results in slowed cellular growth and reduced incorporation of EdU (Fig. 2B). Immunostaining with antibodies specific for TUJ1, TH and DDC 48 hours after induction, revealed changes in morphology with *iHIF2α* expressing cells extending TUJ1^+^ neurite-like protrusions (Fig. 2 D-E) and substantially increased levels of TH, DDC and MAP2 (Fig. 2 F-L).

**Figure 2.**
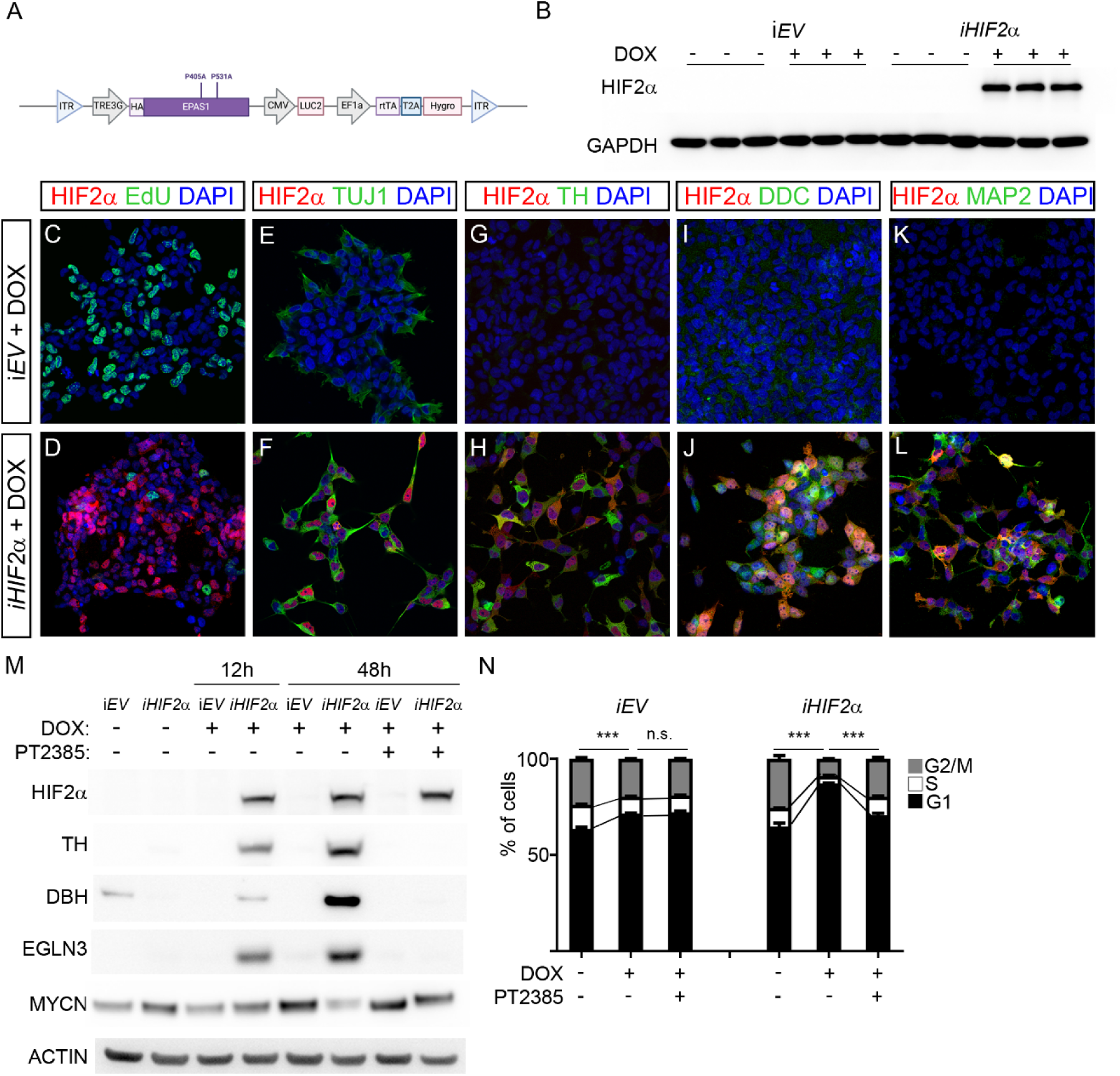
Overexpression of HIF2α in *MYCN*-amplified LAN-1 neuroblastoma cells leads to reduced proliferation and upregulation of chromaffin associated factors. (A) Schematic representation of *piggyBac* construct for doxycycline inducible expression of a stabilized form of *EPAS1* (P450A, P531A). (B) Doxycycline induced expression of *EPAS1* (*iHIF2α)* in LAN-1 neuroblastoma cells. (C-D) Immunostaining with HIF2α (red) combined with EdU (green) and DAPI (blue) in *iEV* control cells (C) and in *iHIF2α* (D) cells treated with doxycycline. (E-F) Immunostaining with HIF2α (red) combined with TUJ1 (green) and DAPI (blue) in *iEV* control cells (E) and in *iHIF2α* (F) cells treated with doxycycline. (G-H) Immunostaining with HIF2α (red) combined with TH (green) and DAPI (blue) in *iEV* control cells (G) and in *iHIF2α* (H) cells treated with doxycycline. (I-J) Immunostaining with HIF2α (red) combined with DDC (green) and DAPI (blue) in *iEV* control cells (I) and in *iHIF2α* (J) cells treated with doxycycline. (K-L) Immunostaining with HIF2α (red) combined with DDC (green) and DAPI (blue) in *iEV* control cells (K) and in *iHIF2α* (L) cells treated with doxycycline. (M) Western blot showing upregulation of HIF2α, TH, DBH and EGLN3 12h after doxycycline induction. Addition of the HIF2α inhibitor PT2385 at 48h reverses increase of TH, DBH and EGLN3, but MYCN protein levels are restored. ACTIN is shown as loading control. (N) Cell cycle analysis after PI-staining shows a decrease in cells in S-phase and G2/M-phase. This decrease is reversed upon treatment with PT2385 in *iHIF2α* cells. Cell cycle data is represented as mean ± SD; P-value of differences in S-phases was calculated with ANOVA with Tukey’s multiple comparisons test.

To further validate that the induction of TH and DBH as well as the reduced proliferation was a response to high HIF2α activity, we used the HIF2α inhibitor PT2385 which inhibits the dimerization of HIF2α with ARNT1 and prevents activation of HIF2α target genes (34–36). Already after 12 hours of doxycycline treatment there was an increase in protein levels of TH, DBH and the HIF2α target EGLN3. After 48 hours this effect was even more pronounced. However, treatment with PT2385 abolished the effect without affecting induced HIF2α levels (Fig. 2M). We could also detect a reduction in MYCN protein levels upon *iHIF2α* induction, this decrease was partially reversed by PT2385 (Fig. 2M). To quantify the effects on cellular proliferation we performed propidium iodine (PI) staining. This revealed a significant reduction in cells in S and G2/M phase upon *iHIF2α* induction which was reversed upon PT2385 treatment (Fig. 2N).

### Overexpression of *EPAS1*/HIF2α in *MYCN*-amplified neuroblastoma cells induces a transcriptional response characterized by genes highly expressed in noradrenergic cells of the adrenal medulla

To investigate the full transcriptional response to HIF2α, we prepared cDNA libraries of three different clones expressing *iHIF2α* for sequencing 12 hours and 96 hours after doxycycline induction. Analysis showed that at 12 hours 2801 genes were significantly upregulated (adj. p<0.05) and 2652 genes were downregulated (Fig. 3A and Supplementary table 1). At 96 hours 1828 genes were upregulated and 1193 were downregulated (Fig. S4A and Supplementary table 1). Gene set enrichment analysis (GSEA) utilizing the “Hallmarks” collection of gene sets from Broad Institute (37, 38), revealed that both at 12 hours and 96 hours induction of HIF2α resulted in the upregulation of several gene sets including the “HYPOXIA” gene set (Fig. 3B-C, S4B-C). In the *iHIF2α* induced cells there was a strong negative correlation with terms associated with MYC-targets and proliferation (Fig. 3B and D and Fig. S4B and D).

**Figure 3.**
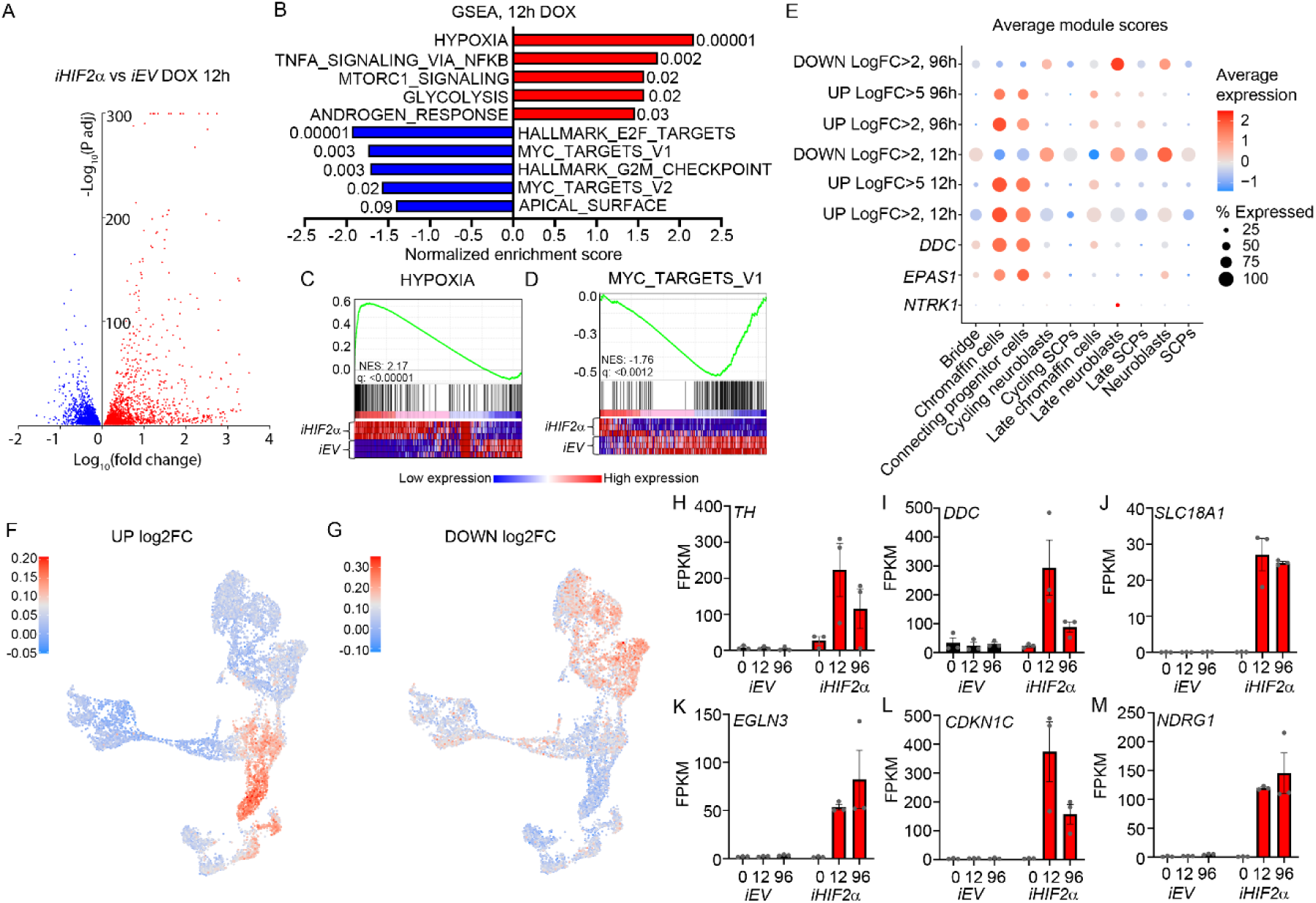
Overexpression of *EPAS1* rapidly induces expression of genes associated with hypoxia and the chromaffin cell lineage while there is a reduction of MYCN targets and genes associated with cell cycle progression. (A) Volcano plot showing genes upregulated (red) and downregulated (blue) 12 hours after induction of *iHIF2α*. (B) Gene set enrichment analysis (GSEA) of the genes from (A), numbers indicate FDR. (C) GSEA of the “HYPOXIA” gene set. (D) GSEA of the “MYC_TARGETS_V1”. (E) Dot plot showing the average gene module scores for the indicated genes and gene groups. (F) Genes upregulated >Log2FC in (A) plotted on the developing adrenal medulla from Jansky et al. (G) Genes downregulated >Log2FC in (A) plotted on the developing adrenal medulla from Jansky et al. (H-M) Expression of the indicated genes *iEV* and *iHIF2α* at 0, 12 and 96 hours after doxycycline treatment.

To understand whether the upregulated or downregulated gene signatures resemble any specific cellular populations in the developing human adrenal medulla, we utilized the Jansky (20) dataset to compute gene module scores of up- and downregulated genes. Both at 12 hours and 96 hours genes upregulated logfold >2 and >5 were enriched in the “Chromaffin cells” and “Connecting progenitor cells” cell types (Fig. 3 E-F, Fig. S4E and Supplementary table 1). In contrast, genes downregulated logfold <2 were enriched for in other cell types, in particular in “Neuroblasts” and “Late neuroblasts” (Fig. 3E and G, S4F and Supplementary table 1). Downregulated genes at both timepoints were depleted in “Chromaffin cells” and “Connecting progenitor cells” (Fig. 3E). To understand the effect of elevated HIF2α levels on the expression of specific noradrenergic chromaffin associated genes, over time, we plotted the FPKM of *TH, DDC* and *SLC18A1* which all showed robust increased expression levels upon *iHIF2α* induction (Fig. 3 H-J). In addition, we plotted expression levels of HIF2α target gene *EGLN3*, the cell cycle inhibitor *CDKN1C* and the tumour suppressor *NDRG1* which is a direct negative target of MYCN regulation (39) (Fig. 3 K-M). There was however no increase in *PNMT* expression.

A recent study from Bishop and colleagues (40) shows how high levels of HIF2α in the mouse adrenal medulla promotes a *Pnmt1^-^/Epas1^+^/Rgs5^+^* noradrenergic phenotype that produces noradrenaline in contrast to the *Pnmt1^+^/Epas1^-^/Rgs5^-^* adrenaline producing cells. In our system the induction of *iHIF2α* induced high levels of *RGS5* (up∼66x, adj. p<1.3e-44) at 96 hours as well as the atypical mitochondrial regulators *COX4I2* (up∼259x, adj. p<8.7e-09) at 12h and *NDUFA4L2* (up∼5x, adj. p<1.2e-04) at 96 hours, which also were upregulated in the mouse adrenal medulla upon induction of HIF2α (40). In addition, analysis of the 498SEQC dataset showed that *RGS5, NDUFA4L2* and *COX4I2* are all significantly correlated with *EPAS1* expression in neuroblastoma and that both *RGS5* and *NDUFA4L2* are profoundly correlated with increased overall survival.

### Induction of *EPAS1*/HIF2α in *MYCN*-amplified neuroblastoma cells rapidly depletes MYCN protein levels, followed by an increase in enzymes necessary for noradrenaline synthesis

Despite a significant negative correlation between *MYCN* and *EPAS1* expression in 498 sequenced clinical samples (6) and a HIF2α dependent reduction in MYCN protein levels (Fig. 2M), we observed no significant increase in *MYCN* expression upon *iHIF2α* induction. The observed downregulation of MYC targets upon *iHIF2α* induction prompted us to investigate how elevated levels of HIF2α affects MYCN protein levels. Following 24 hours of doxycycline treatment there was a clear reduction in MYCN immunoreactivity (Fig. 4A-B). To better map the consequences for MYCN over time we conducted a time-course analysis over 2-96 hours post-doxycycline induction. This revealed that rising levels of HIF2α rapidly reduced MYCN protein levels, followed by an increase in DDC levels (Fig. 4C). Immunostaining at short intervals revealed that there was a window at 2 hours when MYCN and HIF2α proteins were detectable in the same cells. However, already after 4-8 hours, there was a distinct loss of MYCN as HIF2α levels rose (Fig. 4 C-G). The reduction in MYCN protein levels aligns with the gene sets enriched for downregulated genes as indicated by GSEA analysis. Retinoic acid (RA) is used as an adjuvant treatment in patients with high-risk tumours and has a strong differentiation inducing effect in certain neuroblastoma cell lines, however not in LAN-1 cells. In LAN-1 cells the response to *iHIF2α* induction was a depletion of MYCN protein levels and induction of DDC, whereas RA alone failed to promote any such response (Fig. 4I). The combined induction of *iHIF2α* and RA treatment appears to increase the levels of DDC even further, suggesting that high levels of HIF2α might restore RA sensitivity in LAN-1 cells. Treatment with the proteasome inhibitor MG132 restored MYCN protein levels in *iHIF2α* cells treated with doxycycline for 24 hours (Fig. 4J). Thus, high levels of HIF2α results in targeting MYCN for proteasomal degradation and even in the presence of MG132 the levels of MYCN were lower than in control cells (*iEV* + DOX). In contrast, the levels of DDC induced by *iHIF2α* expression, were unaffected by MG132 treatment, indicating that HIF2α induces expression of the *DDC* gene rather than influencing its protein stability (Fig. 4J).

**Figure 4.**
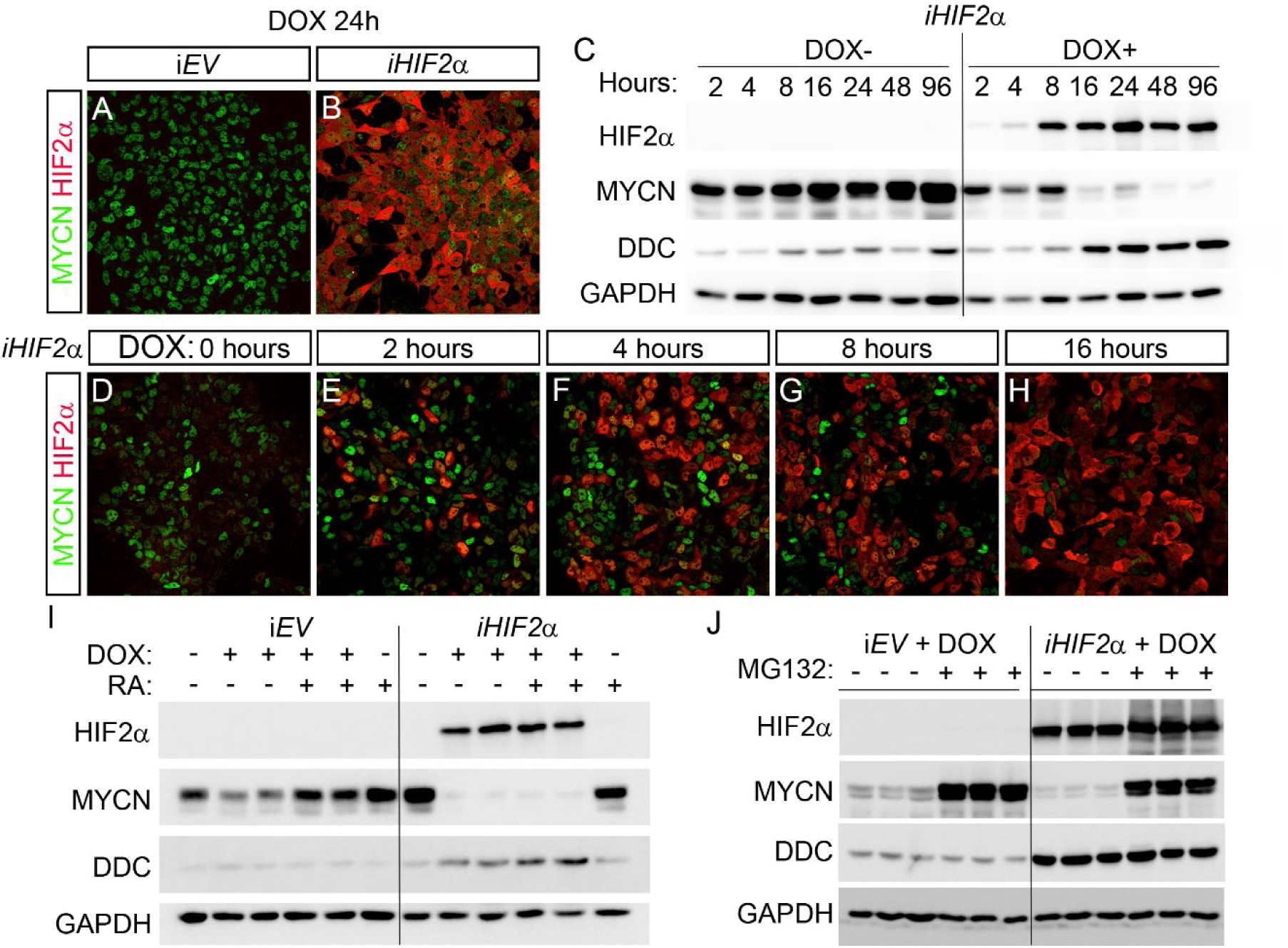
Induction of *iHIF2α* depletes MYCN protein levels prior to the increase of DDC. (A-B) In *iEV* expressing cells there is no induction of HIF2α and MYCN protein levels remain high, 24h after doxycycline treatment (A), whereas in *iHIF2α* cells there is an upregulation of HIF2α and an reduction in MYCN protein levels (B). (C) Western blot showing protein levels of HIF2α, MYCN, DDC and GAPDH at the indicated time points after doxycycline induction in *iHIF2α* cells. (D-H) Immunostaining with antibodies for MYCN (green) and HIF2α (red) at the indicated time points after doxycycline induction. (I) Western blot showing the indicated protein levels in LAN-1 cells treated with retinoic acid (RA) with or without doxycycline induction. (J) Western blot showing the indicated protein levels in LAN-1 cells treated with the proteasome inhibitor MG132 after 24h of doxycycline induction.

To validate the effects of high levels of HIF2α in another *MYCN*-amplified neuroblastoma cell line we established a similar *piggyBac* system in the SK-N-BE(2) neuroblastoma cells. As in the LAN-1 cells doxycycline induced high levels of HIF2α and reduced incorporation of EdU (Fig. S5 A-B). Immunostaining for TUJ1, TH, DDC and MAP2 revealed increased immunoreactivity and changes in cellular morphology similar to the changes in LAN-1 cells (Fig. S5 C-J). As in the LAN-1 cells *iHIF2α* induction resulted in decreased MYCN immunoreactivity (Fig. S5 K-L). We also investigated whether there was cell autonomous loss of MYCN in the cells that upregulate HIF2α. Reflecting the effect in LAN-1 cells, immunostaining for MYCN and HIF2α revealed an inverse correlation in protein levels (Fig. S5 M-Q). This pattern was also supported by western blot analysis reflecting the time dependent decrease in MYCN protein levels and an increase in DDC, TH, and DBH protein levels (Fig. S5R). PI staining showed a significant reduction in the number of cells in S phase upon induction of *iHIF2α* (Fig. S5S).

### In neuroblastoma xenografts, induction of *EPAS1*/HIF2α impedes tumour growth and triggers expression of genes highly expressed in noradrenergic chromaffin cells

The strong reduction of MYCN protein levels upon activation of the HIF2α pathway in *MYCN-* amplified cells coupled to the drop in proliferation and expression of noradrenergic genes (Fig. 4) indicate that rather than acting as a neuroblastoma oncogene, HIF2α has potential tumour suppressive capacity. To test this in an animal model we xenografted nude mice with LAN-1 cells expressing *iHIF2α* or the *iEV* empty vector as a control. Tumours were allowed to grow to 500mm^3^ before treatment with doxycycline was initiated. Although a proportion of cells did not respond to doxycycline, the induction of HIF2α efficiently impeded tumour growth (Fig. 5 A-B). Neuroblastoma cells in the HIF2α-induced tumours, exhibiting high levels of HIF2α also have high levels of TH, but low levels of KI67, whereas control tumours exhibit high levels of KI67 and low levels of TH (Fig. 5 C-F).

**Figure 5.**
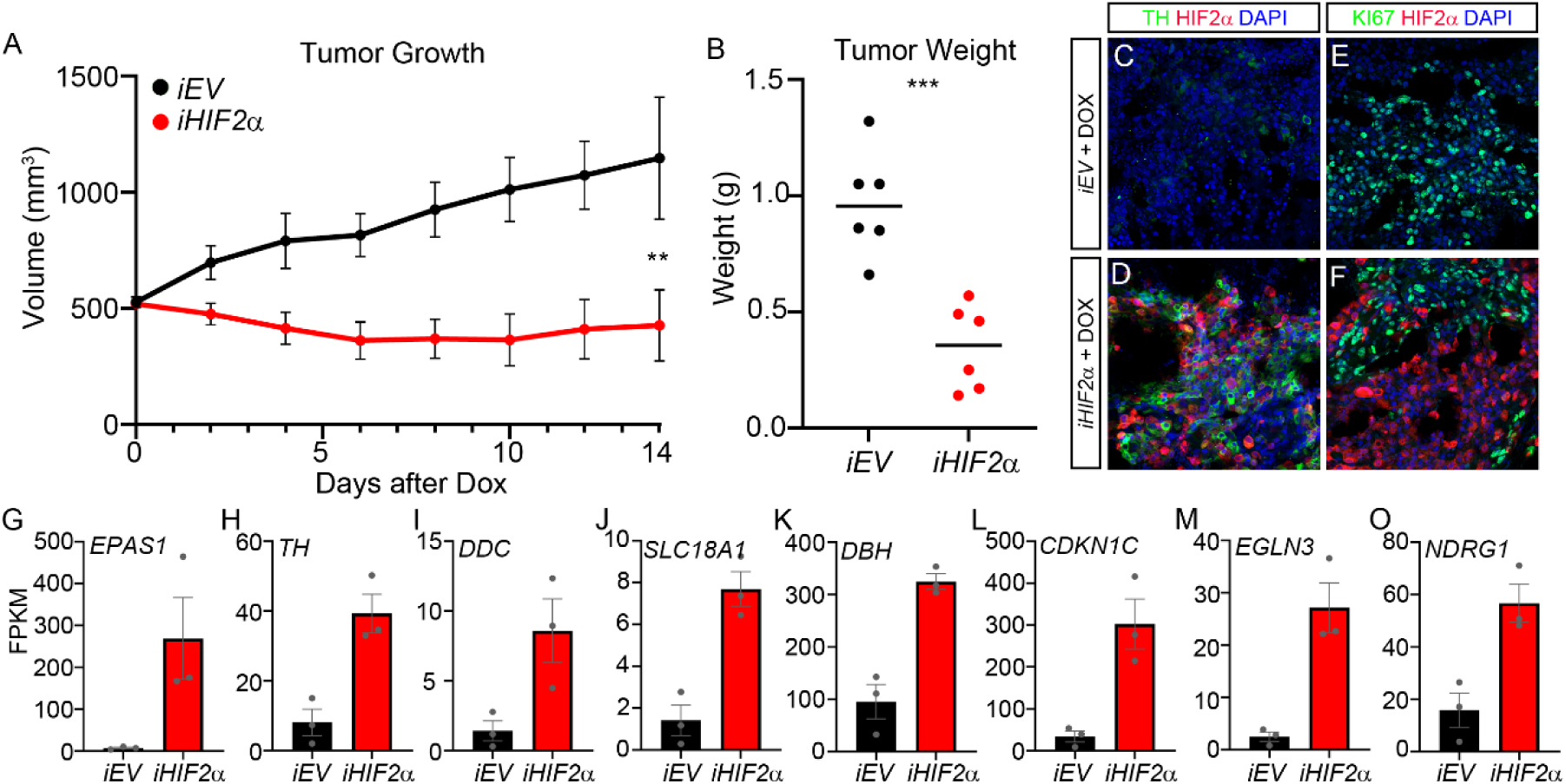
Induction of *iHIF2α* in formed xenograft tumours impedes tumour growth and upregulates expression of chromaffin genes. (A) Induction of *iHIF2α* in xenograft tumors with an average volume of 500mm^3^ significantly impedes tumor growth. (B) Tumor weight at the endpoint for doxycycline treated *iEV* tumors (black) and *iHIF2α* tumors (red). (C-D) Immunostaining for TH (green) and HIF2α (red) in doxycycline treated xenografted tumors, *iEV* (C) and *iHIF2α* (D). (E-F) Immunostaining for KI67 (green) and HIF2α (red) in doxycycline treated xenografted tumors, *iEV* (E) and *iHIF2α* (F). (G-O) Expression of the indicated genes in doxycycline treated xenografts. Tumour growth in A is represented as mean tumour volume ± SD, **p<0.01, two-way ANOVA, (n =6 per group), with Šídák’s multiple comparisons test. In B, ***p<0.001, unpaired t-test with Welch’s correction.

Despite a heterogenous cellular response to doxycycline-mediated induction of *iHIF2a* (Fig. 5 D and F) we performed gene expression analysis on retrieved post treated tumours. Besides high levels of *EPAS1* (Fig. 5G), a similar cohort of genes to those upregulated after 12h and 96h after *in vitr*o induction (Fig. 3H-M) (*TH, DDC, SLC18A1, DBH, CDKN1C, EGLN3* and *NDRG1*) was upregulated in the xenografts wherein *iHIF2α* was induced (Fig. 5 H-O). Thus, *EPAS1* correlated well with expression of markers of noradrenergic cells within the adrenal medulla, e.g. *DBH* or *TH* as well as the cell cycle inhibitor *CDKN1C* and the tumour suppressor *NDRG1* (Fig. 5 G-O). There was however no increase in *PNMT* expression. This supports our previous analysis of *EPAS1* expression within the same lineage during mouse development (7) as well as our analysis of *EPAS1* expression within the development of the human lineage and in neuroblastoma (Fig. 1 and Fig. S1).

## Discussion

Our study reveals that high levels of *EPAS1* is strongly associated with genes highly expressed in noradrenergic cells of the adrenal medulla. This gene repertoire is also predictive of low-risk tumours lacking *MYCN*-amplification with better patient outcome. Furthermore, overexpression of *EPAS1*/HIF2α in *MYCN*-amplified neuroblastoma cells leads to a rapid depletion of MYCN protein, followed by a significant reduction in proliferation rate and induction of a transcriptional response reflecting gene expression in the noradrenergic chromaffin lineage of the developing adrenal medulla. In a xenograft model of neuroblastoma, induction of HIF2α in tumours that had reached an average volume of 500mm^3^, efficiently impeded tumour growth. Analysis of the excised tumours revealed an absence of HIF2α immunoreactivity in certain regions, which coincided with high immunoreactivity for the cell cycle marker KI67 and low TH staining, emphasizing the inverse relationship between high levels of HIF2α and neuroblastoma tumour growth as well as indicating that the suppressive effect of *iHIF2a* on tumour growth could be underestimated.

The reduction in MYCN protein levels occurs rapidly, within 2-4 hours of doxycycline treatment. This suggests that HIF2α may facilitate the targeting of MYCN for proteasomal degradation, which is supported by the relative stabilization of MYCN protein levels upon treatment with the proteasome inhibitor MG132. Given the central role for MYCN and its target genes in high-risk neuroblastoma, the abrupt and potent depletion of MYCN upon HIF2α induction strongly argues against an oncogenic role for HIF2α in this particular disease. The results in this study are consistent with an earlier study wherein dual treatment of neuroblastoma cells with the demethylating agent 5-Aza-deoxycytidine and retinoic acid inhibited xenografted tumour growth while inducing *EPAS1* expression along with several HIF2α target genes (6). Notably, under those conditions there was a distinct reduction in *MYCN* gene expression, together with the partial restoration of MYCN levels upon PT2385 treatment (Fig. 2M), this implies that under certain conditions high levels of HIF2α impinges on the expression of *MYCN.* Despite the rapid reduction of MYCN protein levels, our sequence analysis revealed no reduction in *MYCN* expression upon *iHIF2α* induction after 12 h or 96h. This could either be explained by HIF2α directly affecting MYCN stability or by secondary effects, involving MYCN target genes. Alternative explanations and combinations of effects are also possible, hence the mechanisms through which HIF2α regulates MYCN levels warrant further studies.

The high levels of *EPAS1* expression in the chromaffin lineage during adrenal medulla development potentially reflects a critical role of HIF2α in the development of catecholamine-secreting cells in the sympathoadrenal lineage, more specifically for the noradrenergic N-type rather than adrenergic E-type cells. Interestingly, HIF2α has been shown to be necessary for differentiation of neural crest derived catecholamine producing cells in mice (30) and for differentiation in the vertebrate CNS (41) as well as being able to induce a *Pnmt^-^*/*Epas1^+^*/*Rgs5^+^*noradrenergic phenotype in the mouse adrenal medulla (40). In a recent study from Adameyko and colleagues, the development of the of the chromaffin cell lineage in mouse is described in detail (22). Two end states of differentiation are delineated with the main difference between the two subtypes being the expression of *Pnmt* i.e. E-type cells (*Th*^+^/*Dbh*^+^/*Pnmt^+^*) producing adrenaline and N-type cells (*Th*^+^/*Dbh*^+^/*Pnmt^-^*) producing noradrenaline. In addition, they identify a third terminally differentiated population expressing high levels of *Epas1*, which they hypothesize represents a mature group of oxygen sensing chromaffin cells. To our knowledge, the human chromaffin cell lineage has not been analyzed with the same resolution and there are discrepancies in marker expression between mouse and human chromaffin populations, so if a similar tripartite organization exist in humans is not known. There is some overlap between genes upregulated by *iHIF2α* induction and different genes defining the three of the chromaffin cell groups identified in the study from Adameyko and colleagues, but there is no definite pattern that perfectly fits any of the populations. We don’t fully understand the role of HIF2α for chromaffin differentiation. However, the clear correlation between N-type cells and high levels of *EPAS1* as well as the upregulation of genes highly expressed in N-type cells but not the *PNMT* gene upon *iHIF2α* induction in neuroblastoma cells suggest a role for HIF2α in the generation of noradrenergic chromaffin cells but not adrenergic chromaffin cells.

The reduction in MYCN levels is followed by a rapid upregulation of noradrenergic markers and cell cycle exit, i.e. events reflecting the acquisition of a more mature cellular state. Whether it is the downregulation of MYCN per se that triggers this differentiation or whether lower levels of MYCN creates a transcriptional environment more amenable to signals inducing differentiation remain to be fully determined. However, the oncogenic impact of MYCN in neuroblastoma is hard to underestimate and the profound reduction in MYCN levels is likely a key factor in the cellular response to the induction of *iHIF2α*.

In a recent study on the role of the tumour suppressor gene *VHL*, Kaelin and colleagues show that upon inactivation of *VHL,* the HIF2α protein levels rise (42). This results in substantially reduced cell fitness in neuroendocrine cancer cells and induction of NDRG1, a member of N-MYC downregulated gene family. Importantly this also substantially attenuates tumor formation in xenografts of Kelly neuroblastoma cells. Even though this study focuses on the regulation of HIF2α by VHL in another context, it shows that high levels of HIF2α is not compatible with neuroendocrine tumour cell fitness.

Our study has not addressed all aspects of neuroblastoma and we cannot exclude the possibility that HIF2α can be involved in processes such as metastasis, relapse, drug-resistance or in hypoxia associated events affecting tumour pathophysiology. However, our computational analysis of several neuroblastoma and adrenal medulla data sets combined with functional experiments *in vitro* and in *in vivo* preclinical models of neuroblastoma strongly suggest that HIF2α is not an oncogene in neuroblastoma but rather harbour tumour suppressive characteristics, which is probably connected to the reduction of MYCN protein levels. Despite a validated role as an oncogene in renal cell carcinoma (4), HIF2α exhibits tumour suppressive properties in several other types of cancers (43–50). Given these differences it is feasible that the role of HIF2α is highly context dependent.

Taken together, our results challenge the predominant view that HIF2α is an oncogene or even a facilitator of tumour growth of in neuroblastoma.

## Materials and methods

Neuroblastoma (NB) cell lines LAN-1 and SK-N-BE(2) were cultured at 37°C in a humidified incubator with 95% air and 5% CO2. Cells were routinely maintained in 100 mm culture dishes with RPMI-1640 medium (Gibco, 31870) supplemented with 2 mM L-glutamine (Gibco, 25030), 100 units/mL penicillin, 100 μg/mL streptomycin (Gibco, 15140), and 10% fetal bovine serum (FBS, HyClone, SV30180). Cells were subcultured regularly to maintain exponential growth.

### Western Blot Analysis

Upon completion of Dox treatment, LAN-1 and SK-N-BE2 cells were collected and lysed with ice cold EBC lysis solution (50 mM Tris, pH 8.0, 120 mM NaCl, 0.5% Nonidet P-40). Protein concentration was measured through Bradford assay (Biorad). Equal amounts of proteins were loaded on 10% SDS PAGE, followed by transfer on nitrocellulose membrane at 90volt for 1.5 hours. Following transfer, the membrane was blocked with 5% Skim milk in 1x Phosphate Buffer Saline+0.5% Tween20 (PBST) for 30minutes. After 30 minutes, membranes were blotted with primary antibodies for NMYC (1:500, SC 53993), DDC (1:500, SC 293287), TH (1:1000, PelFreez P40101-150), HIF2α (1:1000, Cell Signal, 7096) dissolved in PBST overnight in cold room. Next day, the membranes were incubated with anti-mouse (Cell Signal, 7076 s) or anti-rabbit (Cell Signal, 7074 s) IgG conjugated secondary antibody solution for 1.5-2 hours and then developed with ECL immobilon western chemiluminescent HRP substrate (Millipore, WBKLS0500). Images were captured by Amersham and LAS4000 imaging systems. Membranes were then reblotted with either GAPDH (1:3000) or β-ACTIN (1:3000) next day for internal normalisation.

### Immunostainings

Following completion of treatment, equal number of cells (grown on coverslips) were fixed with 4% PFA in room temperature for 20minutes. Gradual fixation with 2% PFA for 5 minutes and then 4% PFA for 15 minutes was done for LAN-1 and SH-SY5Y cells which were kept in long-term culture. Following fixation, cells were blocked with 5% BSA, 0.2% Tween 20 and 0.02% Sodium Azide in 1X PBS solution for one hour, followed by overnight antibody incubation in cold condition. Next day, cells were incubated with a combination of DAPI and appropriate Alexa fluorophore conjugated secondary (anti-mouse A2102, anti-Rabbit A31572, Thermofisher scientific) antibody solutions for 90 minutes. Afterwards cover slips were mounted on slides. Images were acquired in Leica SP8 confocal microscopy at 40x zoom, 1024×1024 dimension.

### EdU proliferation assay

The assay was performed according to the Kit protocol (C10637, ThermoFisher Scientific) Briefly, 20µM EdU was added to each experimental point for the last one hour of the total treatment duration (on 23rd hour for 24 hours of Dox treatment). Following an hour of incubation, cells were fixed with 3.5% PFA for 15 minutes followed by permeabilization in 0.5% TritonX-100 in 1X PBS for 20 minutes. Meanwhile, a mixture of reaction buffer, Copper protectant, Alexa Fluor 488, EdU buffer additive was prepared and added to each point after 20 minutes of permeabilization. Upon completion of 30 minutes, reaction mixture was removed. Cells were then washed twice and subjected to general protocol for counterstaining with desired antibodies (HIF2α/HA/MAP2).

### PI staining and cell cycle analysis

For SK-N-BE(2) cells in Fig. S5Q: After 24 hours of Dox treatment, cells were dispersed into single cells with 1X PBS-0,5mM and collected on ice. Cells were then washed with ice cold 1X PBS for three times followed by fixation and immobilization in ice cold 70% ethanol for one hour. After one hour, fixed cells were washed twice with ice cold 1X PBS and then subsequently incubated with a propidium iodide (PI) staining solution containing 100μg/mL RNase A, 0.1 % Triton-X 100 and 10μg/mL PI for 30 min. Stained cells were washed with 1X PBS twice and subjected to flow cytometer (Accuri C6+, BD Biosciences). All experimental samples were prepared in triplicate. Data were collected from 10000 gated cells and analysed further.

For LAN-1 cells in Fig. 2N: Following the respective treatments, cells were harvested by trypsinization and fixed with pre-chilled 70% ethanol (EtOH) at 4°C for 1 hour. After fixation, cells were washed with phosphate-buffered saline (PBS) and incubated on ice for 15 minutes in PBS containing 0.25% Triton X-100 to permeabilize the membranes. Cell pellets were then resuspended in PBS containing 10 μg/mL RNase A and 20 μg/mL propidium iodide (PI) and incubated at room temperature in the dark for 30 minutes. Cell cycle analysis was performed using flow cytometry (BD FACSCanto II). All experimental samples were prepared in triplicate. Flow cytometry data were analyzed using FlowJo software.

### Vectors and stable cell lines construction

The HA-HIF2α-P405A/P531A-pBabe-puro plasmid was generously provided by the Susanne Schlisio laboratory. The PiggyBac transposase vector pCAG T7 K hyPBase was a gift from the Kenneth Chien laboratory. Primers for cloning were designed, and the HA-HIF2α-P405A/P531A construct was inserted into a custom-made inducible PiggyBac Transposon vector, pB-TRE-Luc2. All cloned sequences were verified by Sanger sequencing, performed by Integrated DNA Technologies (IDT).

To generate stable cell lines, the transposon vectors pB-TRE-HA-HIF2α-P405A/P531A-LUC2 or pB-TRE-LUC2, along with the transposase vector pCAG-T7 K hyPBase, were co-transfected into cells using Lipofectamine 2000 (Invitrogen) at a 4:1 vector-to-transposase ratio. Forty-eight hours post-transfection, cells were selected in 200 μg/mL Hygromycin B until non-transfected control cells were completely eliminated. This approach successfully established an inducible, stable HIF2α expression system, functional even under normoxic conditions.

Cells were seeded either directly into culture dishes or on top of coverslips placed in dishes at approximately 50% confluence. The following day, 50 ng/mL doxycycline (Clontech Laboratories) was added to the culture medium, along with either 10 μM of PT2385 (Sigma-Aldrich) or 0.1% DMSO (Sigma-Aldrich) as a control, for the designated treatment duration. The medium containing doxycycline and PT2385 was replenished every 2 days. At the conclusion of the experiment, cells were either fixed with 4% paraformaldehyde (PFA, Sigma-Aldrich) or harvested for subsequent experimental analyses.

### RNA Sequencing and Data Analysis

Total RNA was extracted from cultured cells or xenografted tumor tissues using the RNeasy Kit (Qiagen) according to the manufacturer’s instructions. RNA quality and quantity were assessed using a Bioanalyzer (Agilent Technologies) to ensure high-quality RNA input. RNA sequencing (RNA-seq) libraries were prepared using the TruSeq RNA Library Preparation Kit v2 (Illumina) following the standard protocol. Paired-end RNA sequencing was performed by Novogene, and high-throughput sequencing data were processed for quality control, alignment, and differential expression analysis using a combination of established bioinformatics tools, such as FastQC, STAR, and DESeq2. Gene expression profiles were further analyzed for pathway enrichment and functional annotation.

### Statistical analysis

For the analysis in Fig. 1J-M expression data normalized and standardized was kindly provided by Bedoya-Reina et al. 2021 and correspond to that obtained with the PAGODA pipeline. Plots were done using Scanpy. For significance calculations in Fig. 1M, Welch’s t-test was used. Cell cycle data in Fig. 2N and S5Q were analyzed using ANOVA with Tukey’s multiple comparisons test. Xenograft growth in Fig. 5A was analyzed with two-way ANOVA, with Sídák’s multiple comparisons test and tumour weight in Fig. 5B was analyzed using Unpaired t-test with Welch’s correction.

### Gene Module Scores Analysis

Gene module scores were computed using the *AddModuleScore* function from the Seurat R package, version V5.0.0 (51). The reference dataset for the analysis, encompassing embryonic adrenal medulla data, was obtained from Jansky et al. (<u>available at:</u> https://adrenal.kitz-heidelberg.de/developmental_programs_NB_viz/). To identify genes for module score calculation, differentially expressed genes (DEGs) were filtered based on the absolute log2FoldChange values derived from DEG analysis. Two gene sets were created by applying less stringent (|log2FoldChange| > 2) and high-stringency (|log2FoldChange| > 5) thresholds, respectively. These filtered gene lists were subsequently used as input gene modules for the *AddModuleScore* function in Seurat, generating module scores for each individual cell. The resulting scores were added to the reference dataset as additional metadata.

### Mouse Xenografts

A suspension of 2 × 10^6 neuroblastoma cells in 200 μL of a 1:1 mixture of phosphate-buffered saline (PBS) and Matrigel (BD Biosciences, 354248) was injected subcutaneously into the right flank of adult female Crl (Ncr)-Foxn1^nu^ (nude) mice. Tumor growth was monitored by measuring the external dimensions using a digital caliper every other day. The greatest longitudinal diameter (length) and the greatest transverse diameter (width) were recorded, and tumor volume was calculated using the modified ellipsoid formula: Tumor volume= [(length × width^2^)/2]

Mice were provided with a doxycycline-supplemented diet (0.625 g/kg Doxycycline Hyclate) once the tumors reached an approximate volume of 500 mm^3. The doxycycline diet was replenished every 2 days. Throughout the experiment, overall health status was carefully monitored, and body weight was measured weekly. At the end of the study, mice were euthanized in accordance with approved protocols, and tumors were excised. Tumor weights were measured, and the samples were snap-frozen for subsequent analyses.

### Immunofluorescence Staining on Xenograft Cryosections

Xenograft tumors were excised at the endpoint of the experiment and immediately snap-frozen in optimal cutting temperature (OCT) compound (Sakura Finetek) using liquid nitrogen. Frozen tumors were sectioned into 8–10 μm thick slices using a cryostat (Leica), and the sections were mounted onto Superfrost Plus glass slides (ThermoFisher Scientific). Slides were air-dried at room temperature for 30 minutes before being fixed in cold 4% paraformaldehyde (PFA) for 15 minutes at 4°C.Following fixation, the sections were washed three times in phosphate-buffered saline (PBS) and permeabilized with 0.1% Triton X-100 in PBS for 10 minutes. Non-specific binding was blocked by incubating the sections in PBS containing 5% normal goat serum (NGS) for 1 hour at room temperature. Primary antibodies were diluted in PBS with 1% NGS and applied to the sections, followed by incubation overnight at 4°C in a humidified chamber. After incubation with primary antibodies, the sections were washed three times with PBS and incubated with fluorophore-conjugated secondary antibodies (Invitrogen) together with DAPI (4’,6-diamidino-2-phenylindole, Sigma-Aldrich) for 1 hour at room temperature in the dark. After wash with PBS, slides were mounted with ProLong Gold Antifade Mountant (Thermo Fisher Scientific) and coverslipped. Immunofluorescence images were acquired using a confocal laser scanning microscope (Zeiss, LSM700).

## List of antibodies

| Target | Cat.No. | Supplier | Application | Host |
| --- | --- | --- | --- | --- |
| β-ACTIN | A5441 | Sigma | WB | Mouse |
| **GAPDH** | ab22555 | abcam | WB | Rabbit |
| **HA** | sc-7392 | Santa Cruz | WB, ICC | Mouse |
| **HA** | 3724S | Cell Signaling | WB, ICC, IHC, | Rabbit |
| **HIF2α** | 7096 | Cell Signaling | WB | Rabbit |
| **Ki67** | ab16667 | Abcam | WB, IHC, ICC | Rabbit |
| **MYCN** | sc-791 | Santa Cruz | WB | Rabbit |
| **DBH** | 8586 | Cell Signaling | WB, | Rabbit |
| **TH** | P60101-150 | Pel-Freez Biologicals | WB, IHC, ICC | Sheep |
| **TH** | P40101-150 | Pel-Freez Biologicals | WB, IHC, ICC | Rabbit |

### Ethical considerations

All animal experiments were performed according to Swedish guidelines and regulations, the ethical permit 6420–2018 was granted by ‘Stockholms Norra djurförsöksetiska nämnd, Sweden’. Neuroblastoma primary tumors came from the Swedish neuroblastoma Registry (ethical permission (DN03-736) granted by Karolinska Institutets Forskningsetikommitté Nord (clinical information described in Li *et al.* (52).

Primary neuroblastoma (NB) tumor samples were obtained from the Swedish Neuroblastoma Registry, with ethical clearance (DN03-736) provided by the Regional Ethical Review Board at Karolinska Institutet, Stockholm, Sweden. Prior to sample collection, written informed consent was obtained from the families of all subjects, in compliance with ethical standards and data protection regulations.

## Acknowledgments

Support was provided by The Swedish Research Council (VR 2022-000731 to J.H), The Swedish Children Cancer Foundation, The Swedish Cancer Foundation and Kempefonderna.

