## Supplementary figures for "HIF2α negatively regulates MYCN protein levels and promotes a low-risk noradrenergic phenotype in neuroblastoma"

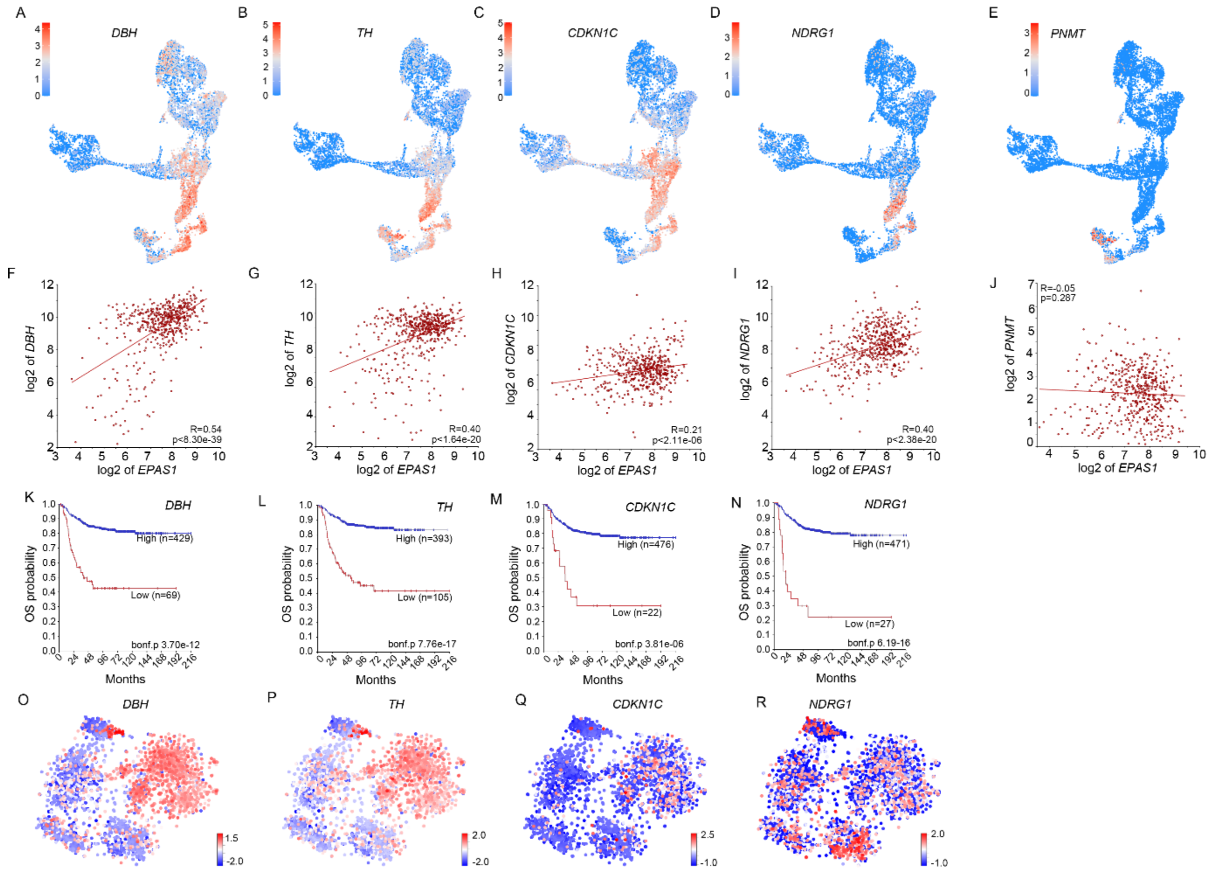


**Supplementary Figure S1. *EPAS1* expression is associated with the chromaffin cell lineage, *cyclin dependent kinase inhibitor 1C,* and the *NDRG1* gene downregulated by MYCN.**

(A-D) *DBH* (A), *TH* (B), *CDKN1C* (C) and *NDRG1* (D) expression are enriched in connecting progenitor and chromaffin cells in the developing human adrenal medulla, dataset from Jansky et al.

(E) *PNMT* expression is enriched in late chromaffin cells.

(F) Expression of *DBH* and *EPAS1* are positively correlated in neuroblastoma tumors.

(G) Expression of *TH* and *EPAS1* are positively correlated in neuroblastoma tumors.

(H) Expression of *CDKN1C* and *EPAS1* are positively correlated in neuroblastoma tumors.

(I) Expression of *NDRG1* and *EPAS1* are positively correlated in neuroblastoma tumors.

(J) Expression of *PNMT* and *EPAS1* are not correlated in neuroblastoma tumors.

(K) *DBH* expression is correlated to increased overall survival.

(L) *TH* expression is correlated to increased overall survival.

(M) *CDKN1C* expression is correlated to increased overall survival.

(N) *NDRG1* expression is correlated to increased overall survival.

(O) Mapping of *DBH* expression in tSNE of neuroblastoma single cell nuclei show enrichment in noradrenergic low-risk tumor cells, dataset from Bedoya-Reina et al.

(P) *TH* expression is enriched for in noradrenergic low-risk tumor cells.

(Q) *CDKN1C* expression is enriched for in noradrenergic low-risk tumor cells.

(R) *NDRG1* expression is enriched for in endothelial cells.

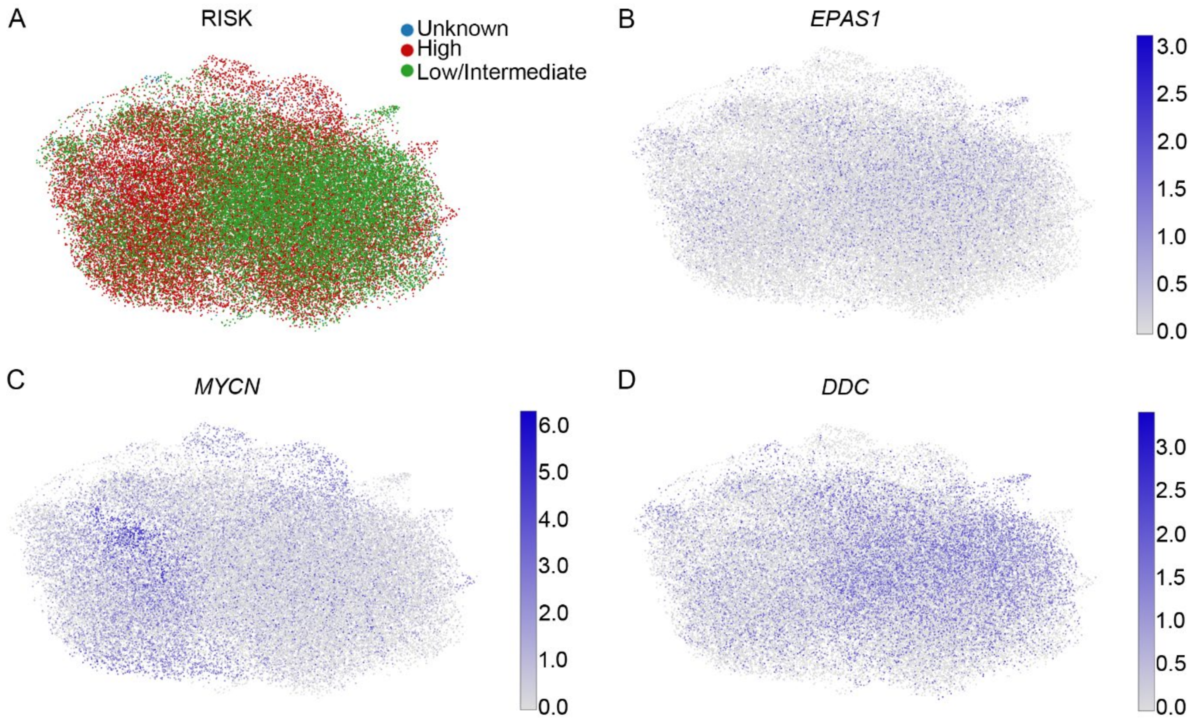


**Supplementary Figure S2. *EPAS1* and *DDC* expression are prominent in low-risk tumor cells with low levels of *MYCN* expression.**

(A) UMAP plot showing neuroblastoma cells of different risk levels, according to Bonine et al.

(B) Expression of *EPAS1*.

(C) Expression of *MYCN*.

(D) Expression of *DDC*.

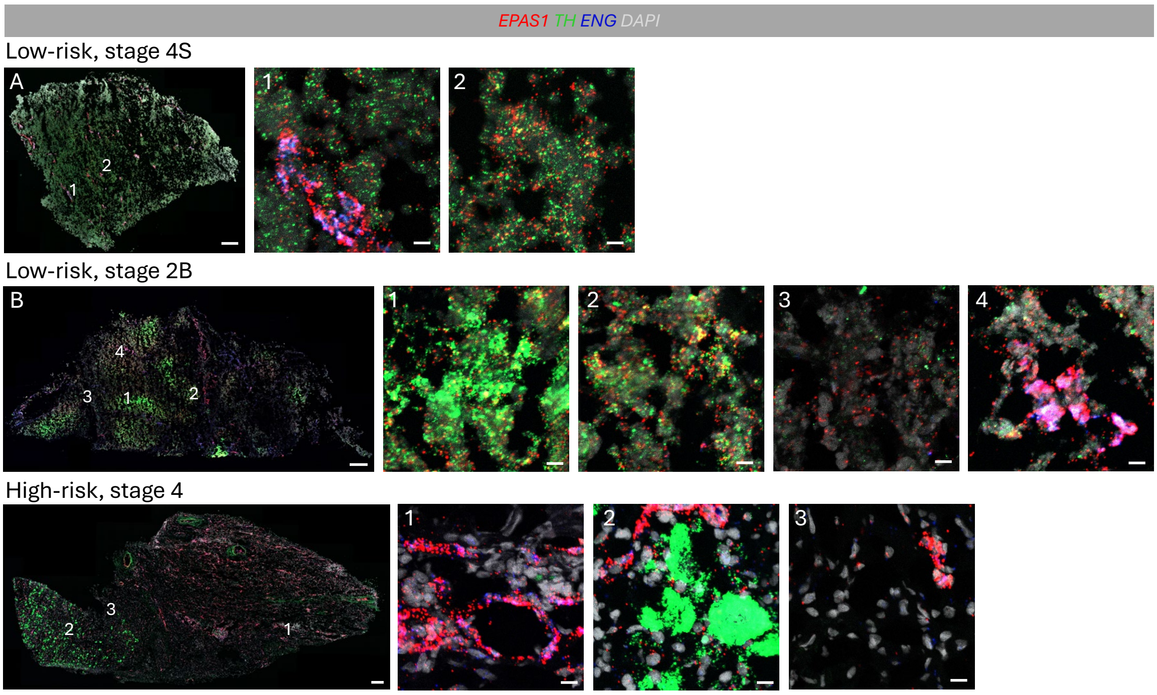


**Supplementary Figure S3. Tiled-scanned 20x images of RNAscope *in situ* hybridization showing *EPAS1, TH* and *ENG* expression in neuroblastoma tumours of different stages.**

RNAscope with probes for *EPAS1* (red), *TH* (green), *ENG* (Blue) and DAPI in white to visualize nuclei.

(A) Low-risk, stage 4S tumour, 1 and 2 represent zoomed in regions as indicated.

(B) Low-risk, stage 2B tumour, 1, 2, 3 and 4 represent zoomed in regions as indicated.

(C) High-risk, stage 4 tumour, 1, 2 and 3 represent zoomed in regions as indicated.

S4

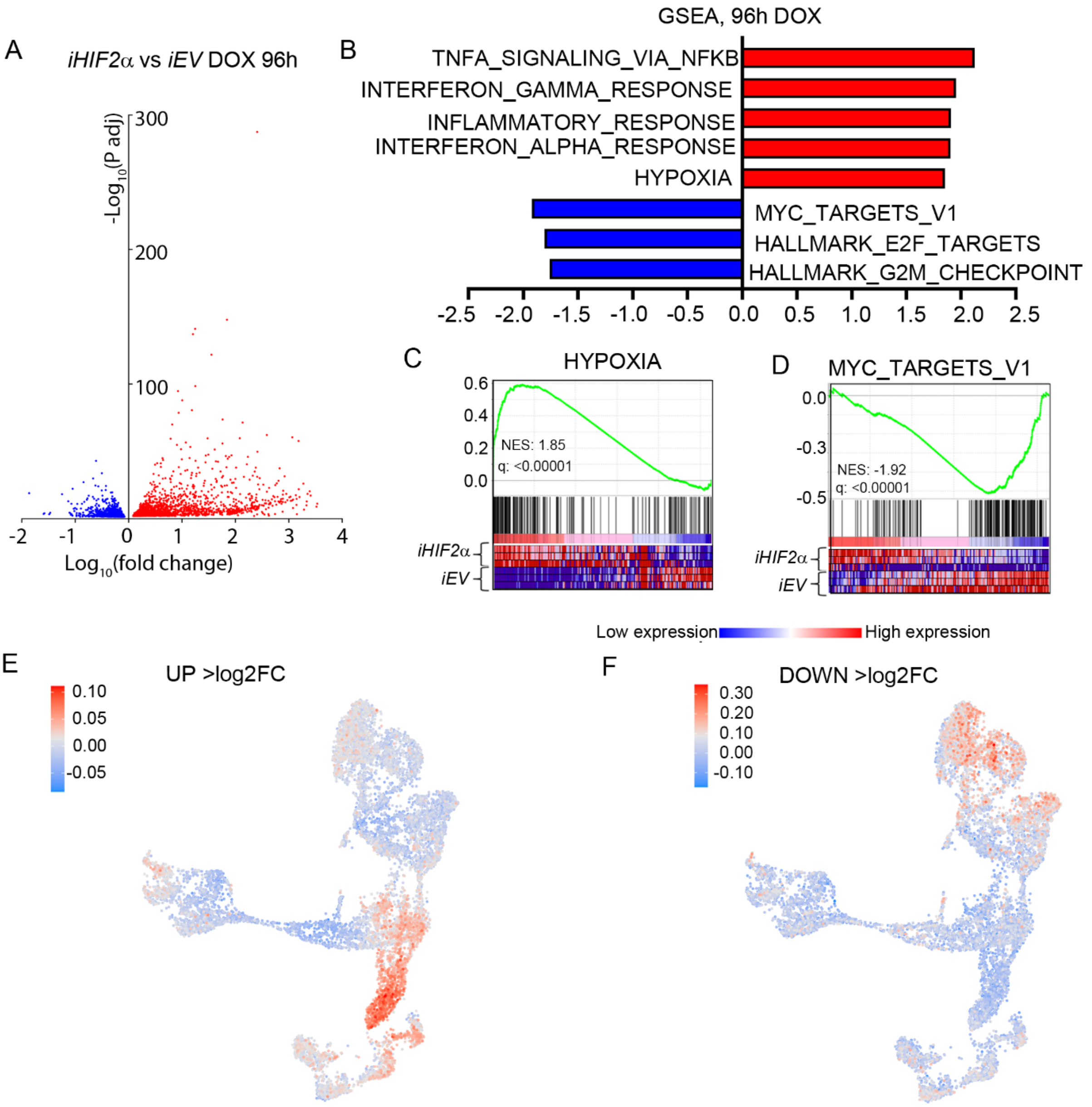


**Supplementary Figure S4. After 96h overexpression of *EPAS1* causes expression of genes associated with hypoxia and the chromaffin cell lineage while there is a reduction of MYCN targets and genes associated with cell cycle progression.**

(A) Volcano plot showing genes upregulated (red) and downregulated (blue) 96 hours after induction of *iHIF2α*.

(B) Gene set enrichment analysis (GSEA) of the genes from (A).

(C) GSEA of the "HYPOXIA" gene set.

(D) GSEA of the "MYC_TARGETS_V1".

(E) Genes upregulated >Log2FC in (A) plotted on the developing adrenal medulla from Jansky et al.

(F) Genes downregulated >Log2FC in (A) plotted on the developing adrenal medulla from Jansky et al.

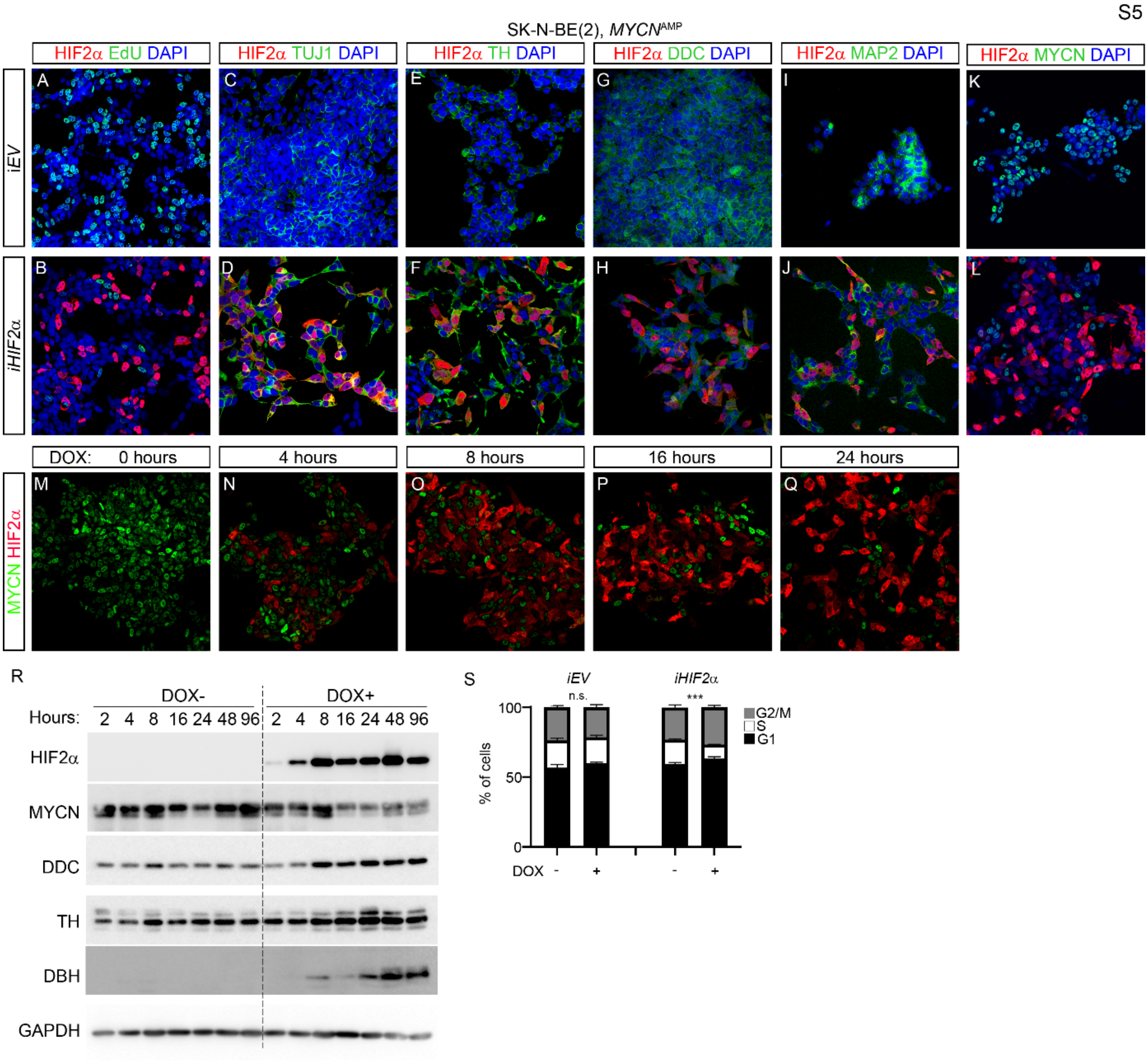


**Supplementary Figure S5. Overexpression of HIF2α in *MYCN*-amplified SK-N-BE(2) neuroblastoma cells leads to reduced MYCN protein levels and upregulation of noradrenergic chromaffin cell associated factors followed by a reduction in proliferation.**

(A-B) Immunostaining with HIF2α (red) combined with EdU (green) and DAPI (blue) in *iEV* control cells (A) and in *iHIF2α* (B) cells treated with doxycycline.

(C-D) Immunostaining with HIF2α (red) combined with TUJ1 (green) and DAPI (blue) in *iEV* control cells (C) and in *iHIF2α* (D) cells treated with doxycycline.

(E-F) Immunostaining with HIF2α (red) combined with TH (green) and DAPI (blue) in *iEV* control cells (E) and in *iHIF2α* (F) cells treated with doxycycline.

(G-H) Immunostaining with HIF2α (red) combined with DDC (green) and DAPI (blue) in *iEV* control cells (G) and in *iHIF2α* (H) cells treated with doxycycline.

(I-J) Immunostaining with HIF2α (red) combined with MAP2 (green) and DAPI (blue) in *iEV* control cells (I) and in *iHIF2α* (J) cells treated with doxycycline.

(I-J) Immunostaining with HIF2$\alpha$ (red) combined with MYCN (green) and DAPI (blue) in *iEV* control cells (I) and in *iHIF2$\alpha$* (J) cells treated with doxycycline.

(M-Q) Immunostaining with antibodies for MYCN (green) and HIF2$\alpha$ (red) at the indicated time points after doxycycline induction.

(R) Western blot showing upregulation of HIF2$\alpha$, DDC, TH, and DBH but downregulation of MYCN after doxycycline induction. GAPDH is shown as loading control.

(S) Cell cycle analysis after PI-staining shows a decrease in cells in S-phase and G2/M-phase upon doxycycline induction of *iHiF2$\alpha$.*

Cell cycle data is represented as mean ± SD; P-value of differences in S-phases was calculated with ANOVA with Tukey's multiple comparisons test.
